# *Daphnia*-associated bacterial communities correlate with diet quantity, environmental conditions and epidemic size across natural outbreaks

**DOI:** 10.1101/2024.06.10.598240

**Authors:** Amruta Rajarajan, Justyna Wolinska, Jean-Claude Walser, Nadine Tardent, Silvana Käser, Esther Keller, Piet Spaak

## Abstract

Zooplankton-associated microbiomes play an important role for host health and contribute to ecosystem processes such as nutrient cycling. Yet, few studies have assessed how environmental gradients and biotic interactions, including parasitism and diet, may shape the microbiome composition of wild zooplankton. Here, we analysed the microbiomes of water fleas from the *Daphnia longispina* species complex using 16S rRNA gene sequencing and a long-term field dataset spanning six sampling events over 13 years. Sampling coincided with outbreaks of the virulent eukaryotic gut parasite *Caullerya mesnili.* Additionally, we explored how microbiome structure varied in relation to water parameters, phytoplankton density (i.e. *Daphnia* diet), and zooplankton density and community structure. *Daphnia* microbiomes displayed strong temporal variation, and comparatively small differences based on host infection status. Microbiome beta diversity correlated with phytoplankton density but not with its community composition, including green algae, protists and cyanobacteria. Environmental conditions, including temperature, dissolved oxygen and cyanobacterial abundance - previously found to drive *Caullerya* epidemics - were also associated with distinct microbiome structures. Importantly, microbiome beta diversity co-varied with infection prevalence, suggesting a link between microbiome shifts, epidemic size, and environmental conditions driving large epidemics. Dominant bacterial taxa correlated with *Daphnia* density, whereas the phylogenetic composition of rare taxa was associated with total zooplankton density. These findings demonstrate the dynamic nature of *Daphnia* microbiomes and suggest potential mechanisms by which they may mediate disease dynamics, particularly through associations with diet quantity, temperature, and host population density.

## Introduction

Zooplankton constitute an essential component of aquatic ecosystems, serving as both consumers and prey. Their interactions with predators, phytoplankton and parasites can shape aquatic ecosystem functioning (Bakhtiyar et al., 2020). Among these interactions, parasites represent an often overlooked but critical functional group, influencing ecosystem processes either directly through biomass generation for consumption, or indirectly by altering host population dynamics and traits (Preston et al., 2016). For instance, zooplankton-parasite associations may play a role in nutrient cycling within aquatic ecosystems (Cáceres et al., 2014; Frost et al., 2008).

Zooplankters also harbour communities of diverse, symbiotic microbes that vastly outnumber free-living microbes in the water column (Tang, 2005). For instance, copepods selectively enrich specific bacterial taxa from their environment, which then disperse back into the water, shaping bacterioplankton community composition (Shoemaker et al., 2019). Bacterial epibionts associated with water fleas (*Daphnia*) contribute to nutrient cycling and the transfer of dissolved organic carbon within the pelagic food web (Eckert & Pernthaler, 2014). Zooplankton-associated bacteria can also help their copepod hosts in digesting toxic cyanobacteria (Gorokhova et al., 2021), and serve as a mechanism of adaptation to toxic diets in other zooplankters such as *Daphnia* (Macke et al., 2017). These general symbiotic relationships between zooplankton and bacteria can exert cascading effects on ecosystem health and functioning, underscoring the importance of characterizing the abiotic and biotic factors such as diet and parasitism, that potentially shape the diversity and composition of bacterial communities.

Host-associated bacterial communities (or microbiomes) influence the health and performance of various aquatic organisms (reviewed in Infante-Villamil et al., 2021). In addition to host traits, parasite traits and environmental conditions, microbiomes are proposed as the fourth vertex of a disease pyramid (Bernardo-Cravo et al., 2020). However, unravelling the forces that potentially influence microbiomes in natural settings is challenging due to their temporal variability and the complex interplay of deterministic (e.g. abiotic environmental gradients, biotic interactions of host species) and stochastic (e.g. horizontal transmission) processes (Sze et al., 2020). Wild host microbiomes can fluctuate over various timescales, spanning from hours to days within a single host generation (Caughman et al., 2021; Minich et al., 2020) to longer timescales of months (Krajacich et al., 2018; Pierce & Ward, 2019). The relatively long-term temporal variation in host microbiomes may be linked to seasonal environmental changes (Sharp et al., 2017; Zhu et al., 2022) or occur independently of seasonal patterns (Epstein et al., 2019). Environmental factors such as temperature and dissolved oxygen influence the composition, diversity and function of aquatic host-associated microbiomes (reviewed in Sehnal et al., 2021). While diet is a strong predictor of microbiome composition in many fish species (Escalas et al., 2021), this relationship does not apply to other aquatic organisms, such as fire salamanders (Wang et al., 2021) or marine zooplankton (Xu et al., 2024).

Additionally, horizontal transmission plays a major role in shaping host microbiomes in aquatic environments (Russell, 2019), with host population density influencing microbial dispersal (Miller et al., 2018).

Despite growing interest in host-microbiome interactions, few studies have investigated microbiome changes in wild host populations over time, particularly in relation to natural parasite epidemics. The microbiome of the reef-building coral *Montastraea faveolata* undergoes strong abundance shifts in response to seasonal variation and the onset of Yellow Band Disease (Kimes et al., 2013). In the fish host *Scomber japonicus,* both the microbiome composition and abundance of a pathogen *Photobacterium damselae* correlated with the summer conditions (Minich et al., 2020). Similarly, amphibian declines attributed to the disease chytridiomycosis are linked to seasonal changes in skin microbiota (Schmeller et al., 2022), with the abundance of specific bacterial taxa correlating with chytrid infection intensity in the frog *Philoria loveridgei* (López et al., 2017). These examples illustrate how host microbiomes may both influence and respond to disease dynamics in natural populations (Stencel, 2021).

The water flea *Daphnia* is a globally prevalent zooplankton genus with significant ecological roles, including controlling phytoplankton populations (Hébert et al., 2016) and influencing bacterioplankton abundance (De Corte et al., 2023) and community composition (Berga et al., 2015; Degans & De Meester, 2002). Conversely, *Daphnia-*associated bacterial communities are also shaped by the surrounding bacterioplankton (Eckert et al., 2021; Macke et al., 2020) and are crucial for host survival and reproduction (Sison-Mangus et al., 2015). The presence of specific bacteria such as *Limnohabitans* sp. in the *Daphnia* microbiome affects host abundance (Peerakietkhajorn et al., 2015)*. Daphnia* microbiome compositions also fluctuate across multiple timescales: influenced by diurnal feeding rates (Pfenning-Butterworth et al., 2022), host age in laboratory conditions (Callens et al., 2018), and seasonal variations in wild populations (Hegg et al., 2021). Moreover, *Daphnia* populations experience strong selection from various microparasites, which can regulate their population densities (Ebert, 2005) and subsequently, their microbiome compositions. For instance, *Daphnia* infected with the virulent eukaryotic gut parasite *Caullerya mesnili* (Class: Ichthyosporea; hereafter *Caullerya*) experience reduced diversity and altered community composition of their associated bacteria (Rajarajan et al., 2022).

Despite advancements in understanding host-microbiome and host-parasite interactions, few studies have characterized their relative effects on wild zooplankton communities, or their association with long-term disease dynamics. Here, we investigate *Daphnia* microbiome variation in Lake Greifensee across six *Caullerya* epidemics spanning 13 years. *Caullerya* commonly parasitizes *Daphnia* guts and is widespread in European lakes (Wolinska et al., 2007). This parasite reduces host fecundity and life span (Lohr et al., 2010), thereby exerting strong selection pressure on host populations (González-Tortuero et al., 2016; Turko et al., 2018). We analyse how *Daphnia* microbiome compositions correlate with *Caullerya* epidemic size, host density and environmental conditions (temperature, dissolved oxygen and abundance of cyanobacteria in lake water), which have been identified as key drivers of *Caullerya* epidemics in this lake (Tellenbach et al., 2016). We also test if the community composition of *Daphnia-*associated bacteria correlates with the composition and density of dominant pelagic phytoplankton (cyanobacteria, green algae, protists, golden algae and diatoms), and zooplankton (copepods, cladocerans, rotifers). We hypothesize that (a) *Daphnia* microbiome composition (i.e. dominant bacterial orders in *Daphnia*) and beta diversity differ between years and by host infection status; (b) environmental gradients (temperature, dissolved oxygen and cyanobacterial abundance), known to drive *Caullerya* epidemics, correlate with *Daphnia* microbiome structure and (c) phytoplankton community composition predicts *Daphnia* microbiome structure, reflecting the role of diet in shaping host-associated microbial communities.

## Material & Methods

### *Daphnia* sampling and processing

*Daphnia* were collected from Lake Greifensee, Switzerland (N 47°20′41″, E 8°40′21″), using vertical towing with a 150 µm zooplankton net from 0-30 m depths at three sampling stations. Zooplankton samples from all stations were pooled into a single 10 L canister half-filled with surface lake water. Samples were collected during the peaks of natural *Caullerya* epidemics on six occasions over 13 years (10/10/2007, 11/08/2011, 29/08/2013, 04/09/2014, 24/08/2017, 10/11/2020), as part of a long-term *Daphnia* infection study (Turko et al., 2018). For clarity, these sampling events are hereafter referred to as “years”, as they represent different calendar years around epidemic peaks (Fig. S1). Upon collection, zooplankton samples were sieved through a ∼250 µm mesh. Adult specimens (>1 mm in size, independent of infection status) belonging to the *Daphnia longispina* species complex were gently lifted by the antennae and screened for *Caullerya* spores in the gut under 50× magnification (Lohr et al., 2010). Individuals with *Caullerya* spores were classified as ‘infected’, whereas those without visible spores were categorized as ‘uninfected’. *Daphnia* with resting eggs (ephippia), or with other infections or rare co-infections involving *Caullerya* and other parasites, were excluded. Individual *Daphnia* were then transferred to wells containing 20 µL autoclaved MilliQ water and stored at -80 °C within eight hours of collection from the lake, until further processing.

### Bacterial community profiling of *Daphnia*

DNA was extracted from whole *Daphnia* individuals using the HotSHOT method (Montero-Pau et al., 2008). Samples were incubated in an alkaline lysis solution at 95 °C for 30 min, cooled on ice for 3-4 min, and eluted in a neutralizing solution. DNA extraction was carried out in three blocks (Table S1). Three DNA extraction negative controls containing only extraction reagents were added in the third DNA extraction block. Extracted DNA was stored at -20 °C until further processing. Subsequent PCR steps were carried out simultaneously for all samples. Bacterial DNA (V4-V5 region of the 16S gene) was amplified using barcoded universal 16S primers 515F (5’-GTGCCAGCMGCCGCGGTAA-3’) (Caporaso et al., 2011) and m806R (5’-GGACTACNVGGGTWTCTAAT-3’) (Apprill et al., 2015) under the following cycling conditions: initial denaturation at 95 °C – 3 min; (98 °C – 20 s, 52 °C – 15 s, 72 °C – 15 s) 33 cycles; followed by a final extension at 72 °C for 5 min. This PCR step was done with three technical replicates for each sample, and 10 PCR no-template negative controls were included. Technical replicates were pooled before the next PCR step. For the second PCR, indexes from the Illumina Nextera XT V2 Library Prep Kit were then added to the samples under the conditions: initial denaturation at 95 °C – 3 min; (95 °C – 30 s, 55 °C – 30 s, 72 °C – 30 s) 10 cycles; followed by a final extension at 72 °C for 5 min. The location of DNA samples (including a total of 13 negative controls, i.e. 3 “blank” DNA extraction and 10 PCR no template samples) in the 384-well plate was randomized throughout the workflow, though technical replicates in the first PCR step were in adjacent wells. Indexed PCR products were then purified using magnetic Agencourt AMPure XP beads (Cat #A63881), quantified using a Qubit HS Assay, normalized and pooled into a 1.2 nM library for amplicon sequencing. Paired-end sequencing was carried out using MiSeq with 600 cycles (PE300) v3 run kit and 10% PhiX. Final sample sizes for analyses (Table S1) were determined after bioinformatic processing (see below).

### Pre-processing of reads

Sequencing generated 7.6M paired-end reads, after exclusion of one infected sample from 2014 that had an unusually high number of reads (∼720,000 as opposed to the average of 46,230 reads in biological samples). Initial pre-processing steps were performed on the Euler computing cluster at ETH Zurich, as previously described (Rajarajan et al., 2022). Raw reads were 5’-end trimmed, merged, and quality-filtered. Amplicons were denoised with UNOISE3 (Edgar & Flyvbjerg, 2015) and the resulting Zero-radius Operational Taxonomic Units (ZOTUs) were assigned taxonomy using the non-Bayesian SINTAX classifier v.138 (Edgar, 2016) against the Silva database v138 (Quast et al., 2013) with a confidence cut-off of 0.75. Biological samples had 320 to 237,358 reads and negative controls had 412 to 2188 reads, with one PCR no-template control unexpectedly showing 24,356 reads. Based on this distribution of reads in biological samples versus negative controls, biological samples with <3000 reads were excluded from downstream analyses. This resulted in a final sample size of 9-16 *Daphnia* per year and infection status, totalling 71 uninfected and 90 infected individuals (Table S1).

Subsequent filtering steps were performed in R v4.0.2 (R Core Team, 2023) using the phyloseq (McMurdie and Holmes 2013) and vegan (Dixon, 2003) packages. 62 ZOTUs (21 of unidentified phyla, 18 chloroplasts, 6 mitochondria, 16 singletons and 1 eukaryote) were filtered out. An additional 42 ZOTUs were removed after rarefying samples to 3,017 reads using vegan::*rarefy_even_depth*, resulting in a final dataset of 423 ZOTUs. There was no consistent increase or decrease in ZOTU richness from older to newer samples, indicating that long-term freezing did not impact microbiome composition (Fig. S2).

### Environmental data for Lake Greifensee

Sampling dates for physicochemical, zooplankton, and phytoplankton data are listed in Table S1. Depth-resolved physicochemical data for Lake Greifensee were obtained from the Office of Waste, Water, Energy and Air of the Canton of Zurich (AWEL, http://www.awel.zh.ch/). Temperature and dissolved oxygen measurements were averaged for the top 15 m of the lake. Phytoplankton were sampled from 0-20 m depth using a Schröder bottle. Phytoplankton (including cyanobacterial) cell counts in lake water were measured with the Utermöhl method (Utermöhl, 1958) using sedimentation chambers. Dominant phytoplankton and microbial groups - diatoms (Fig S3), golden algae (Fig. S4), green algae (Fig. S5), protists (Fig. S6) and cyanobacteria (Fig. S7) were identified to the lowest possible taxonomic level based on morphology, using inverted light microscopy. Zooplankton samples were collected with three vertical tows of a 95 µm plankton mesh at the same locations where *Daphnia* samples were taken, and were preserved in 100% ethanol. To assess *Caullerya* infection rates, 200 adult *Daphnia* were visually screened for *Caullerya* spores in their gut (see above for details).

### Statistical analyses

Statistical analyses were performed in R v4.0.2 (R Core Team, 2023) using the phyloseq (McMurdie & Holmes, 2013) and vegan (Dixon, 2003) packages. Variation in beta diversity (Unifrac distance and Bray-Curtis Dissimilarity) by year and infection status was tested with two-way PERMANOVAs (9999 perm) using vegan::*adonis.* The proportion of variation explained by year and infection status was quantified with a db-RDA using the function vegan::*capscale.* Given the substantial interannual variation in bacterial communities (see Results), subsequent analyses of infection status-effects were carried out separately for each year. Dominant bacterial orders were compared by infection status, where dominant orders were defined as those comprising more than 1% of the dataset or present in more than 50% of all samples. Orders not meeting these criteria were lumped into the category “Other”. Differential abundance of bacterial orders by infection status was tested using the DESeq2 package, with *p*-values adjusted for multiple comparisons using the “fdr” method (Love et al., 2014). Beta diversity in *Daphnia* bacterial communities was correlated with epidemic size (i.e. population infection rates), temperature, dissolved oxygen, and cyanobacterial cell counts using vegan::*envfit*, with *p*-values adjusted for multiple comparisons using the Bonferroni method. To test for potential associations between *Daphnia* bacterial communities and plankton composition in the lake, Jaccard dissimilarity matrices of *Daphnia* bacterial communities were correlated with those of (a) phytoplankton communities and (b) zooplankton communities, using a Mantel test.

## Results

### Bacterial community beta diversity differs between years and marginally by *Caullerya-*infection status

Analyses of beta diversity (Bray-Curtis Dissimilarity and Unifrac distance) revealed that the primary driver of variation in bacterial communities was the sampling year, accounting for about 30% of the observed variation (Table 1). In contrast, infection status accounted for only 1% of the variation. The beta diversity of dominant taxa in *Daphnia*-associated bacterial communities, assessed by Bray-Curtis Dissimilarity (which weighs bacterial taxa by their relative abundances), differed significantly between years and marginally between infected and uninfected *Daphnia*. When phylogenetic relatedness between ZOTUs was considered using Unifrac distance, both year and infection status had significant effects on beta diversity (Fig. 1, Table 1). However, there was no significant interaction between year and infection status in shaping overall microbiome community structure as assessed with beta diversity.

**Figure 1.**
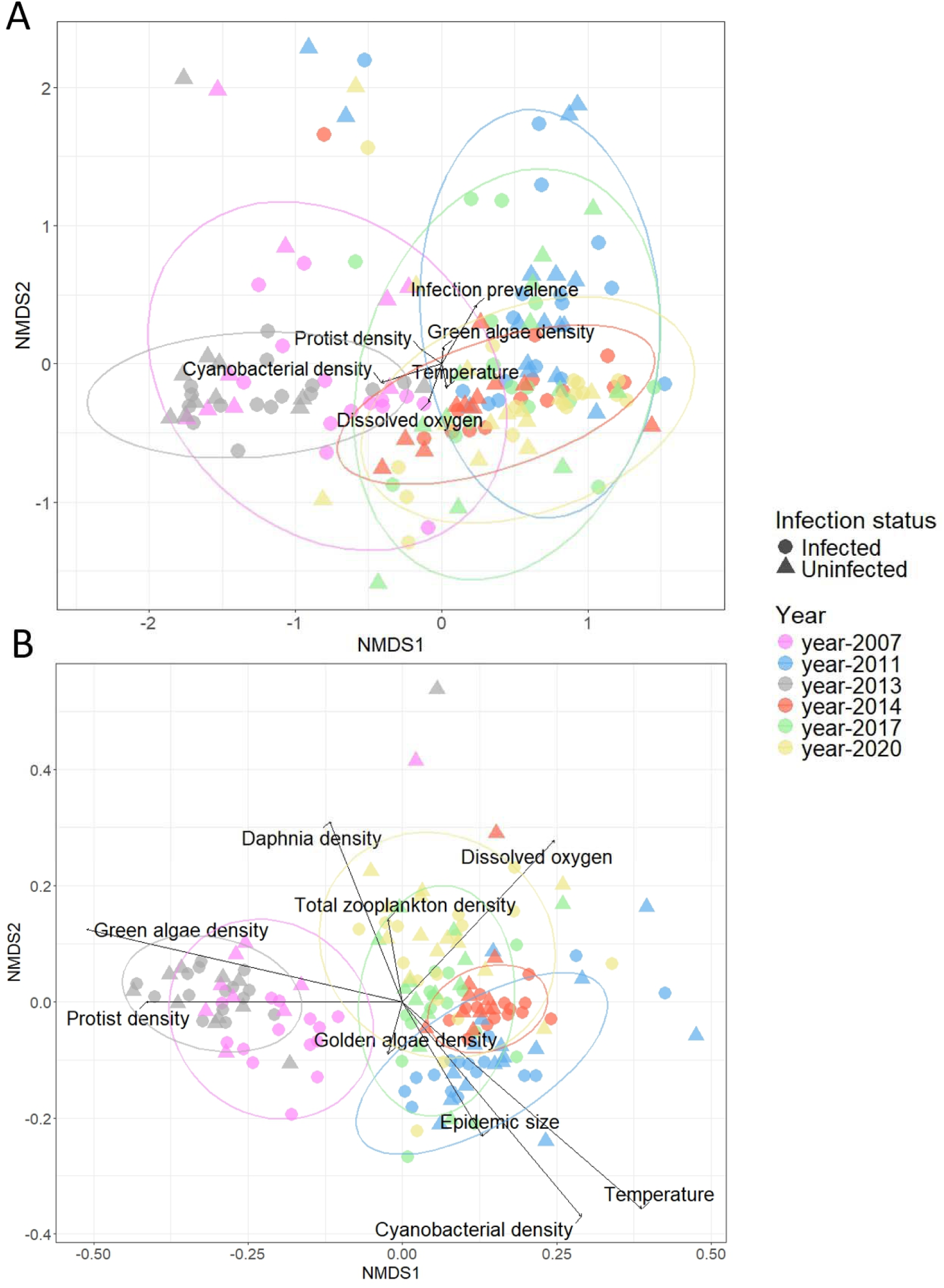
Nonmetric Multi-Dimensional Scaling (NMDS) plots of beta diversity (A) Bray-Curtis Dissimilarity and (B) Unifrac distance in individual *Daphnia*, grouped by sampling year and *Caullerya* infection status. Data point shapes indicate infection status, while colors indicate the year of sampling. Arrows show abiotic (temperature, dissolved oxygen) and biotic (*Caullerya* epidemic size, zooplankton, and phytoplankton densities) variables that significantly correlate with beta diversity. The length of arrows reflects the strength of correlation, and are scaled to the respective NMDS1 and NMDS2 axes, which differ for the two diversity metrics. Ellipses show 95% confidence intervals for each year.

**Table 1.**
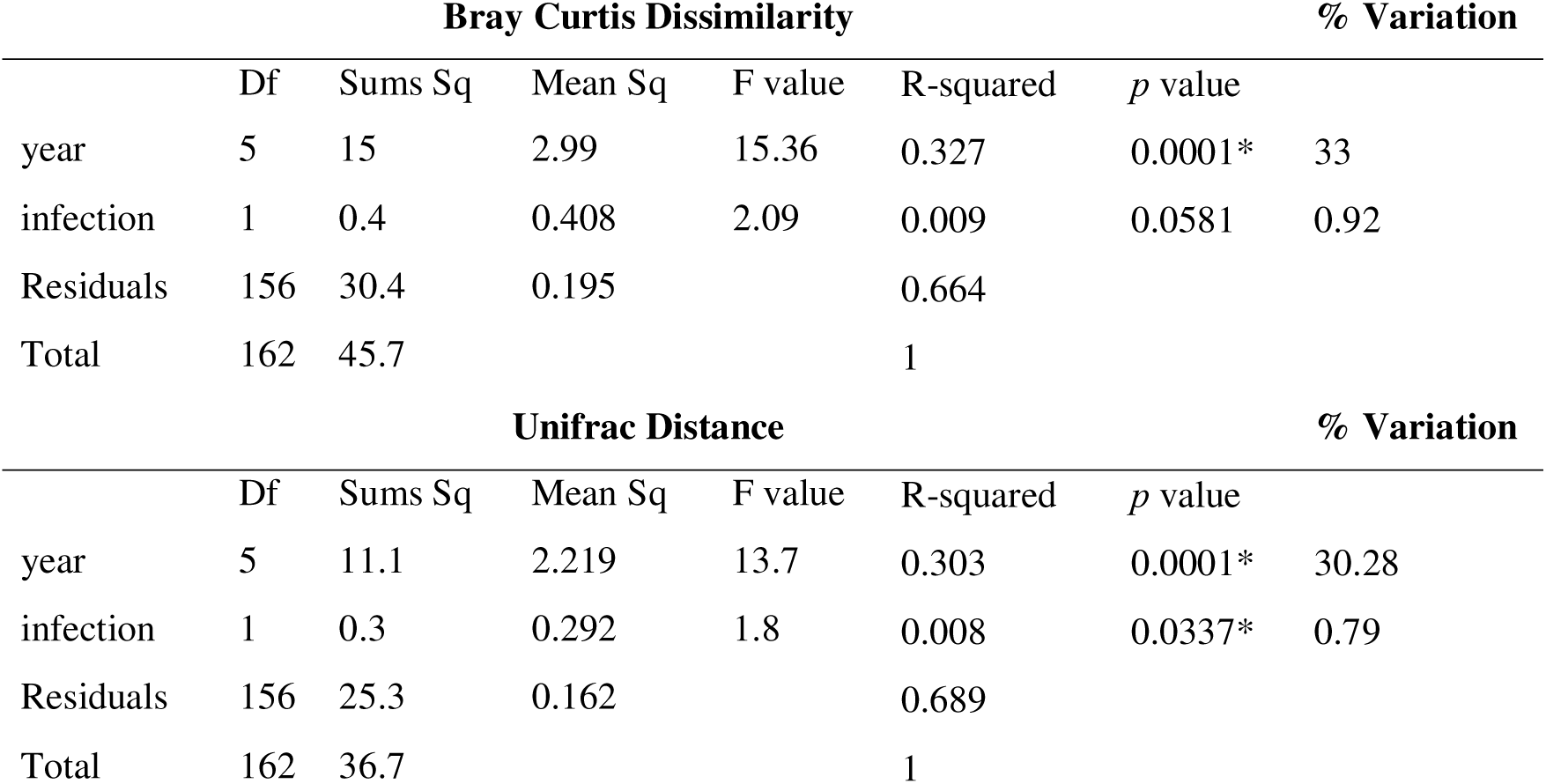
Two-way PERMANOVAs using the beta diversity indices Bray-Curtis dissimilarity (top section) and Unifrac distance (lower section) between years and by and infection status (9999 permutations). Significant results (\**p* < 0.05) are indicated with an asterisk. The “% Variation” column indicates the percentage of variation explained by each factor, as determined by db-RDA. Significant variation in beta diversity by year (but not infection status) may be due differential dispersion of bacterial communities among years, as indicated by vegan::betadisper.

### Relative abundances of dominant bacterial orders differ between *Caullerya-*infected and uninfected Daphnia

Bacterial communities of individual *Daphnia* comprised of 16 dominant orders across all sampling years, with substantial variation among individuals sampled within the same year (Fig. 2). The most abundant bacterial order across all *Daphnia* individuals was Burkholderiales (51.29 ± 24.94%, mean ± SD), followed by Rickettsiales (15.1 ± 26.22%) and Flavobacteriales (14.58 ± 12.73%). Differences in the relative abundances of dominant bacterial orders between *Caullerya-*infected and uninfected *Daphnia* occurred during the epidemic years 2007, 2013 and 2017. In 2007, Caulobacterales was more abundant in uninfected *Daphnia* (2.94 ± 2.36%) compared to infected ones (0.96 ± 0.7%). Conversely, Chitinophagales was more abundant in infected *Daphnia* in both 2007 (1.34 ± 1.22%) and 2013 (2.51 ± 3.97%), compared to uninfected ones (0.45 ± 0.46% and 1.5 ± 2.9%, respectively). In 2013, Rickettsiales was more abundant in infected *Daphnia* (5.18 ± 10.28%) than in uninfected ones (2.12 ± 4.7%). In 2017, Aeromonadales was more abundant in uninfected *Daphnia* (4.64 ± 13.25%) compared to infected individuals (1.66 ± 3.22%), while Pirellulales showed higher abundance in infected *Daphnia* (0.38 ± 0.59%) compared to uninfected ones (0.07 ± 0.12%, Table 2).

**Figure 2.**
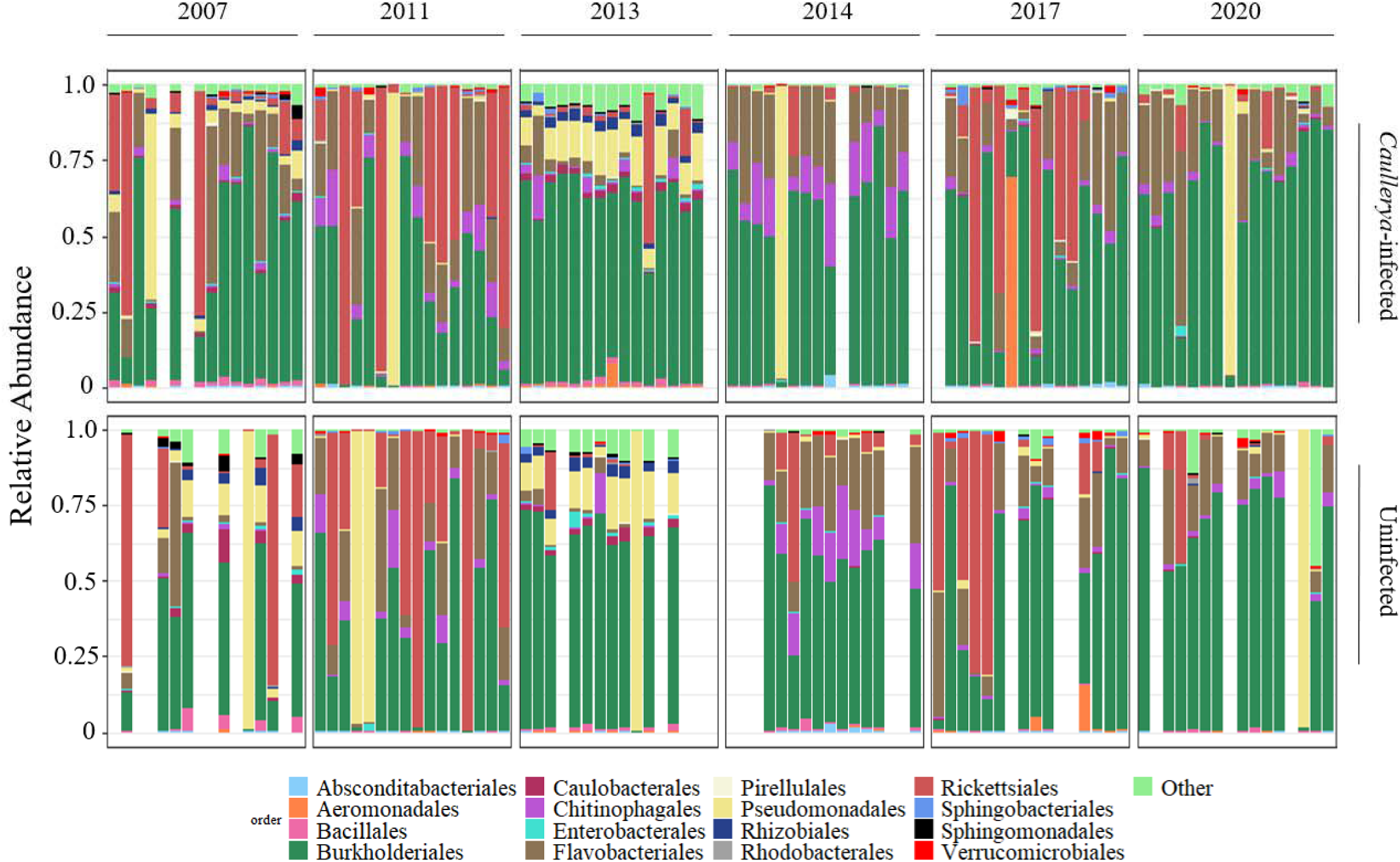
Relative abundances of dominant bacterial orders in individual *Daphnia* across years, with *Caullerya*-infected *Daphnia* in the top row and uninfected *Daphnia* in the bottom row. Each vertical bar corresponds to the bacterial community composition of a single *Daphnia* individual. The category “Other” include bacterial orders comprising <1% of the dataset and present in <50% of samples. Missing bars indicate samples excluded from analyses (see Methods).

**Table 2.** Wald test results comparing the relative abundances of dominant bacterial orders between *Caullerya-*infected and uninfected *Daphnia*, analyzed separately for each year. *p*-values were adjusted for multiple comparisons using the “fdr” method, with significant values (*p*<0.05) shown in bold. The “log2Change” column indicates the logarithmic fold-change in taxon abundance, where negative numbers indicate higher abundance in infected *Daphnia*, and positive numbers indicate higher abundance in uninfected *Daphnia*. The absolute “log2Change” values reflect the magnitude of the difference.

|  | 2007 |  | 2011 |  | 2013 |  | 2014 |  | 2017 |  | 2020 |  |
| --- | --- | --- | --- | --- | --- | --- | --- | --- | --- | --- | --- | --- |
| **Bacterial Orders** | log2Change | *p* value | log2Change | *p* value | log2Change | *p* value | log2Change | *p* value | log2Change | *p* value | log2Change | *p* value |
| Absconditabacteriales | -1.96 | 0.12 | -0.96 | 0.48 | 1.48 | 0.97 | 0.28 | 0.80 | -1.55 | 0.17 | -0.94 | 0.71 |
| Aeromonadales | -1.11 | 0.38 | -1.31 | 0.37 | -0.53 | 0.97 | 1.84 | 0.25 | **3.54** | **0.02** | -1.20 | 0.75 |
| Bacillales | 0.03 | 0.99 | 0.38 | 0.84 | -0.01 | 1.00 | 0.88 | 0.25 | 0.93 | 0.10 | 0.32 | 0.75 |
| Burkholderiales | -1.02 | 0.08 | 0.76 | 0.37 | 0.00 | 1.00 | -0.57 | 0.25 | 0.06 | 0.92 | -0.08 | 0.94 |
| Caulobacteriales | **1.03** | **0.04** | 0.52 | 0.96 | 0.03 | 1.00 | 0.06 | 0.98 | 0.27 | 0.68 | -0.48 | 0.71 |
| Chitinophagales | **-2.04** | **0.04** | -0.09 | 0.97 | **-3.09** | **0.00** | -0.42 | 0.44 | -0.20 | 0.68 | 0.64 | 0.57 |
| Enterobacterales | 0.08 | 0.94 | -1.67 | 0.37 | 0.47 | 0.97 | 1.45 | 0.25 | 0.97 | 0.17 | 1.04 | 0.57 |
| Flavobacteriales | -1.07 | 0.38 | 0.30 | 0.84 | -0.14 | 1.00 | -0.40 | 0.33 | -0.44 | 0.62 | -0.61 | 0.57 |
| Pirellulales | -0.02 | 0.99 | -0.54 | 0.81 | 1.74 | 0.63 | 0.64 | 0.75 | **-2.91** | **0.02** | -1.95 | 0.57 |
| Pseudomonadales | 0.29 | 0.38 | -0.52 | 0.51 | -0.02 | 1.00 | 0.89 | 0.19 | 1.20 | 0.09 | -0.60 | 0.57 |
| Rhizobiales | 0.32 | 0.55 | 0.13 | 0.97 | 0.07 | 0.98 | 0.96 | 0.75 | 0.46 | 0.62 | -0.05 | 0.96 |
| Rhodobacterales | 0.48 | 0.61 | 0.18 | 0.97 | -0.25 | 0.98 | 2.05 | 0.49 | -1.50 | 0.34 | -0.68 | 0.71 |
| Rickettsiales | 1.98 | 0.12 | 0.38 | 0.93 | **-1.88** | **0.05** | 1.64 | 0.19 | -0.25 | 0.92 | 1.03 | 0.57 |
| Sphingobacteriales | -2.11 | 0.08 | 1.08 | 0.37 | -0.60 | 0.97 | -2.64 | 0.19 | -0.41 | 0.68 | 0.65 | 0.71 |
| Sphingomonadales | 1.30 | 0.12 | 0.11 | 0.97 | 0.11 | 0.98 | -0.29 | 0.89 | -0.04 | 0.92 | 0.04 | 0.96 |
| Verrucomicrobiales | -0.82 | 0.55 | -0.89 | 0.41 | -0.68 | 0.98 | 0.92 | 0.44 | 0.88 | 0.34 | 1.22 | 0.57 |
| Other | 0.28 | 0.51 | -0.60 | 0.37 | 0.11 | 0.97 | 0.52 | 0.42 | -0.47 | 0.55 | -0.49 | 0.57 |

### *Daphnia* bacterial community composition correlates with environmental conditions and plankton density

Both Bray-Curtis dissimilarity and Unifrac distance metrics of *Daphnia* bacterial communities showed significant correlations with temperature, dissolved oxygen content, and *Caullerya* epidemic size. Moreover, these metrics were significantly associated with *Daphnia* density and the densities of several phytoplankton and microbial groups in the lake water, such as cyanobacteria, green algae and protists (Fig 1, Table 3). In contrast, only Unifrac distance of *Daphnia* bacterial communities was significantly correlated with total zooplankton density and the density of golden algae in the lake water.

**Table 3.** Correlation of *Daphnia* bacterial community beta diversity indices (Bray-Curtis dissimilarity and Unifrac distance) with abiotic (temperature, dissolved oxygen) and biotic factors (*Caullerya* epidemic size, and densities of phytoplankton and zooplankton groups in lake water), analyzed using vegan::*envfit* (9999 permutations). *p-*values were adjusted for multiple comparisons using the Bonferroni method. Significant results (\**p* < 0.05) are indicated with an asterisk.

| Bray-Curtis dissimilarity |  |  |
| --- | --- | --- |
| Variable | r-squared | p-adjusted |
| Dissolved oxygen (mg/L) | 0.2069 | 0.001* |
| Green algal density | 0.1649 | 0.001* |
| Temperature (°C) | 0.1325 | 0.001* |
| Protist density | 0.0756 | 0.029* |
| Golden algal density | 0.061 | 0.376 |
| Diatom density | 0.0566 | 0.696 |
| Epidemic size | 0.0456 | 0.011* |
| *Daphnia* density | 0.0436 | 0.001* |
| Cyanobacterial density | 0.0378 | 0.001* |
| Total zooplankton density | 0.0089 | 1 |
| Unifrac distance |  |  |
| Variable | r-squared | p-adjusted |
| Temperature (°C) | 0.2776 | 0.001* |
| Green algal density | 0.2771 | 0.001* |
| Cyanobacterial density | 0.2209 | 0.001* |
| Protist density | 0.1718 | 0.001* |
| Dissolved oxygen (mg/L) | 0.1376 | 0.001* |
| *Daphnia* density | 0.1102 | 0.001* |
| Epidemic size | 0.0705 | 0.001* |
| Diatom density | 0.023 | 1 |
| Total zooplankton density | 0.0208 | 0.001* |
| Golden algal density | 0.0087 | 0.001* |

### *Daphnia* bacterial community composition does not correlate with diet (phytoplankton) or zooplankton community composition

Jaccard dissimilarity matrices of *Daphnia* bacterial communities did not exhibit significant correlations with those of phytoplankton, including green algae, golden algae, protists, cyanobacteria, or diatoms (*p* > 0.05, Mantel test). Similarly, *Daphnia* bacterial community composition showed no significant correlation with the composition of pelagic zooplankton, including copepods, rotifers, and *Daphnia* (Table 4).

**Table 4.**
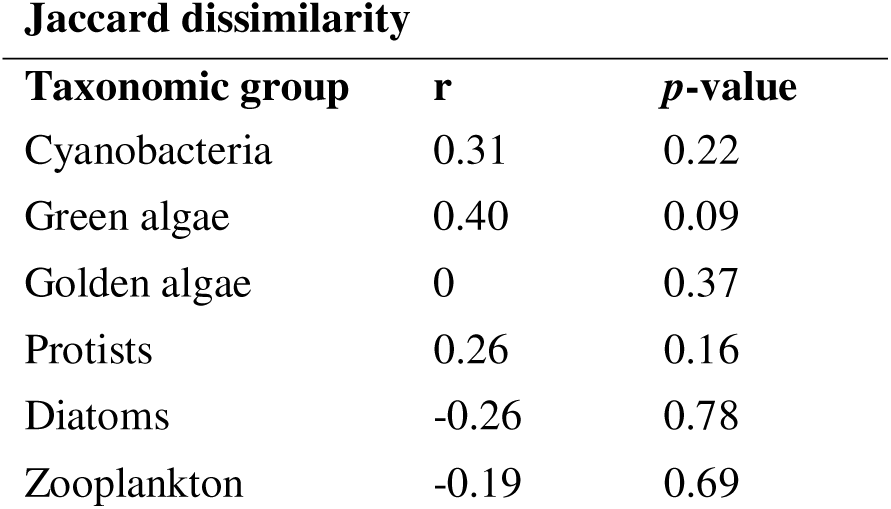
Mantel tests assessing the correlation between presence-absence beta diversity (Jaccard dissimilarity) of *Daphnia-*associated bacteria and phytoplankton taxa (representing *Daphnia* diet) in lake water, across years. The ZOTUs of individual *Daphnia* bacterial communities were pooled for each year. Bacterial community composition of dominant taxa in lake water was determined using microscopy.

## Discussion

We compared the bacterial community (microbiome) composition of wild *Caullerya-*infected and uninfected *Daphnia* collected across six natural epidemics spanning a remarkable 13-year period. This study is the first to investigate potential biotic (parasitism, diet) and abiotic drivers of wild aquatic animal microbiomes over such an extensive timeframe. As with all observational field investigations, the relationships between *Daphnia*-associated bacterial diversity and its potential environmental drivers remain correlational. Nevertheless, many of the investigated factors correlated significantly with microbiome composition, highlighting the dynamic nature of *Daphnia* microbiomes in response to environmental conditions.

We found that *Daphnia* microbiomes differ significantly between years, both in the relative abundances of dominant taxa and their phylogenetic composition. Temporal differences in microbiome beta diversity were more pronounced than differences between infected and uninfected *Daphnia*. While infection status significantly influenced the phylogenetic composition of rare bacterial taxa (Unifrac distance), its effect on dominant taxa (Bray-Curtis dissimilarity) was marginal. Certain bacterial orders such as Chitinophagales exhibited infection-related changes in relative abundance, but this pattern was inconsistent across years. Specifically, infection-associated shifts in dominant bacterial orders were evident in only three of the six surveyed epidemics (2007, 2013 and 2017), with the specific bacterial orders affected varying annually. However, Burkholderiales and Flavobacteriales taxa dominated *Daphnia* microbiomes in all surveyed years. Environmental conditions previously shown to drive *Caullerya* epidemics - including cyanobacterial abundance and water temperature (Tellenbach et al., 2016), as well as *Caullerya* epidemic size itself, correlated significantly with *Daphnia* microbiome compositions. These findings provide the first evidence of a link between aquatic host microbiomes and epidemic size over a long-term, natural study. Our results generally align with studies in other aquatic animals, such as the frog *Philoria loveridgei* (López et al., 2017) and the sponge *Porites astreoides* (Meyer et al., 2014), where temporal variation in host microbiomes was observed, with infection-associated taxa differing by year. Lastly, contrary to our hypothesis, the composition of the *Daphnia* diet (phytoplankton community) did not associate with *Daphnia* microbiome composition, although the abundance of certain phytoplankton groups (green algae and cyanobacteria, but not diatoms) correlated with microbiome structure (beta diversity).

*Caullerya* epidemics are associated with temporal shifts in *Daphnia* population-genetic structure (Schoebel et al., 2011; Turko et al., 2018). Given that *Daphnia* genotypes can harbour distinct microbiomes (Frankel-Bricker et al., 2020; Rajarajan et al., 2023), our findings suggest complex interactions between host genetics, infection status and microbiome compositions. For instance, the order Rickettsiales was enriched in infected *Daphnia* in 2013 but not in other years, including 2020 in the present study. In contrast, we previously noted a putative increase in the relative abundance of Rickettsiales in uninfected compared to *Caullerya-*infected *Daphnia,* among animals collected at a different timepoint during the *Caullerya* epidemic in 2020 (Rajarajan et al., 2022). Similarly, we found no differences in the relative abundance of Enterobacterales between infected and uninfected animals in the present study, including November 2020 (see Fig. S1), whereas Enterobacterales was enriched in infected *Daphnia* from December 2020 (Rajarajan et al., 2022). These patterns suggest that *Daphnia* microbiomes may shift over the course of a single epidemic, consistent with the strong temporal variation detected in our study. Such shifts might be potentially be related to host clonal turnover during epidemics (Turko et al., 2018).

Our results suggest a link between environmental conditions driving *Caullerya* epidemics (temperature, cyanobacterial density) and *Daphnia* microbiomes. Temperature influences *Caullerya* infection prevalence under experimental settings (Schoebel et al., 2011), and natural epidemics are associated with both temperature and high cyanobacterial abundance in lake water (Tellenbach et al., 2016). Moreover, *Daphnia* are more susceptible to *Caullerya* infection when fed a diet consisting of cyanobacteria (regardless of toxicity), as confirmed with a laboratory experiment (Tellenbach et al., 2016). These factors (temperature, cyanobacterial diet) influence not only *Caullerya* infection, but also the composition of *Daphnia* microbiomes (Akbar et al., 2021; Frankel-Bricker et al., 2020; Sullam et al., 2018). In the present study, temperature and cyanobacterial density in the lake water correlated significantly with microbiome composition during *Caullerya* epidemics, with the additional correlation with epidemic size itself suggesting that microbiomes generally differed between *Daphnia* from small versus large epidemics. *Daphnia* microbiomes can mediate tolerance to toxic cyanobacteria (Macke et al., 2017), potentially promoting grazer fitness. This raises the possibility of a bottom-up effect whereby *Daphnia* microbiomes shape host traits that in turn influence phytoplankton abundance. However, this hypothesis requires empirical validation (Decaestecker et al., 2024). The observed correlation between microbiome structure, temperature, and cyanobacterial abundance thus supports the idea that host microbiomes were associated with the specific environmental conditions (temperature and cyanobacterial abundance) that additionally drive disease epidemic size.

Beyond cyanobacteria, the abundance of certain phytoplankton groups (green algae and protists), which are grazed on by *Daphnia,* associated with distinct *Daphnia* microbiome compositions. In natural settings, *Daphnia-*associated bacterial communities vary independently of the composition of bacterioplankton communities (Hegg et al., 2021), whereas laboratory investigations have shown how diet quality shapes their taxonomic composition (Akbar et al., 2021) and functional activity (Li et al., 2021). However, our data did not support the idea that similarity in diet composition (algal and protist communities in lake water) predicts host microbiome composition. This result was contrary to our expectations, as well as reports in marine zooplankton (Xu et al., 2024). Nevertheless, high densities of green algae and protists were associated with distinct *Daphnia* microbiome structures in 2007 and 2013 (see Fig. 1 & Fig. S2). In contrast, the density of diatoms (e.g. *Asterionella* sp.*, Fragillaria* sp. and *Melosira* sp., see Fig. S3) and golden algae (e.g. *Erkenia* sp. (=*Chrysochromulina* sp.), *Uroglena* sp. and *Dinobryon* sp., see Fig. S4) showed no relationship with *Daphnia* microbiome structure, possibly due to their large size, colonial form or motility, making them less ingestible for smaller *Daphnia* species (Cáceres, 1998; Talling, 2003; Wagner, 2008; Yoshida et al., 2001) (albeit golden algal density weakly correlated with the phylogenetic composition of rare bacterial taxa within the *Daphnia* microbiome). High densities of green algae trigger increased *Daphnia* ingestion rates (Porter et al., 1982) which in turn may drive changes in microbiome compositions (Pfenning-Butterworth et al., 2022). Our results posit that food abundance, rather than its specific dietary composition, may play a key role in shaping microbiome structures in wild *Daphnia* populations, consistent with findings that low food availability reduces microbial richness in *Daphnia* (Freese & Schink, 2011; Ichige & Urabe, 2023).

*Daphnia* population density also correlated with the microbiome beta diversity, with microbiomes consistently dominated by Burkholderiales. Burkholderiales taxa are widespread in aquatic hosts including sponges (Paix et al., 2024), corals (Ziegler et al., 2019), and fish (Pérez-Pascual et al., 2021), and are facilitated by ‘symbiotic islands’ or genomic regions promoting colonization and survival within hosts (Stillson et al., 2022). Beneficial Burkholderiales strains may enhance *Daphnia* fitness, potentially driving high host abundances in natural environments (Peerakietkhajorn et al., 2015). Alternatively, high *Daphnia* population densities may facilitate horizontal transmission of microbiota (Miller et al., 2018). Both freshwater (Eckert et al., 2021; Rajarajan et al., 2023) and marine (Boscaro et al., 2022) zooplankton microbiomes are strongly shaped by their environment, with no evidence of phylosymbiosis - i.e. microbiome compositions are not determined by host species identity or relatedness. Horizontal transmission between individuals may therefore be a key factor shaping the microbiomes of wild zooplankton (Russell, 2019) and is known to shape *Daphnia* microbiomes in laboratory conditions (Cooper et al., 2022). Thus, high host population densities may facilitate horizontal transmission of symbionts, increasing their prevalence in hosts. In addition to *Daphnia* population density, we found that the phylogenetic composition of rare, but not dominant, bacteria in wild *Daphnia* microbiomes was correlated with the total density of other zooplankton species in the lake water. This suggests that the presence and abundance of other species in a host community may also influence the structure of wild host microbiomes, although to a lesser extent.

In conclusion, our study highlights the strong temporal variation in *Daphnia* bacterial community composition. Diet quantity, rather than composition, and host population density emerge as key factors potentially shaping these communities. Additionally, epidemic size and environmental conditions that drive disease outbreaks, such as temperature and cyanobacterial abundance, were also associated with microbiome structure. Thus, wild *Daphnia* microbiomes may be linked to host health and disease dynamics through complex interactions involving temperature, diet quantity, and population density.

## Acknowledgement

We would like to thank both past and present members of the Spaak group at Eawag for their experimental support as well as collecting long-term plankton sample data from Greifensee. We also thank the Genetic Diversity Center (GDC) at ETH Zürich for extensive support with 16S rRNA library preparation and sequencing. Special thanks go to Cansu Çetin and Kristel Fernanda Sánchez for stimulating discussions during manuscript preparation, and to Eva Cereghetti for her helpful feedback. We also thank two anonymous reviewers for constructive comments on the manuscript, and Nedim Tüzün for extensive, collaborative brainstorming during manuscript revisions. This work was funded by Eawag (Swiss Federal Institute of Aquatic Science and Technology) and a joint “lead agency” grant from the German Science Foundation (WO 1587/6-1 to JW) and Swiss National Science Foundation (310030 L 166628 to PS).

## Author Contribution

**Amruta Rajarajan**: Conceptualization (equal); Data curation: (lead); Formal analysis (lead); Investigation (lead); Writing – original draft preparation (lead); Writing – review & editing (equal). **Justyna Wolinska**: Conceptualization (equal); Formal analysis (supporting); Writing – original draft preparation (supporting); Writing – review & editing (equal); Funding acquisition (equal); Supervision (equal). **Jean-Claude Walser**: Data curation (supporting); Formal analysis (supporting); Methodology (equal); Writing – review & editing (supporting). **Nadine Tardent**: Investigation (supporting); Writing – review and editing (supporting). **Silvana Käser**: Investigation (supporting); Methodology (equal); Writing – review and editing (supporting). **Esther Keller**: Investigation (supporting); Methodology (equal); Writing – review and editing (supporting). **Piet Spaak**: Conceptualization (equal); Formal analysis (supporting); Writing – original draft preparation (supporting); Writing – review and editing (supporting); Funding acquisition (equal); Supervision (equal).

## Data availability statement

Raw 16S rRNA gene sequences of *Daphnia* samples included in the final analysis are available in the Sequence Read Archive (SRA) under BioProject ID PRJNA1252098. Long-term phytoplankton and zooplankton community data and processed 16S rRNA gene sequence data analysed in this manuscript, as well as R code are available at https://doi.org/10.25678/0008GD.

## Notes

### Competing Interest Statement

The authors have declared no competing interest.

### Summary of Updates

The manuscript including title has been revised to only include correlational language, accurately reflecting the correlational nature of the data. Biological meaning has been derived from these correlations based on a more detailed description of phytoplankton diet.

## References

Akbar, S., Huang, J., Zhou, Q., Gu, L., Sun, Y., Zhang, L., Lyu, K., & Yang, Z. (2021). Elevated temperature and toxic *Microcystis* reduce *Daphnia fitness* and modulate gut microbiota. Environmental Pollution, 271. 10.1016/j.envpol.2020.116409

Apprill, A., Mcnally, S., Parsons, R., & Weber, L. (2015). Minor revision to V4 region SSU rRNA 806R gene primer greatly increases detection of SAR11 bacterioplankton. Aquatic Microbial Ecology, 75(2), 129–137. 10.3354/ame01753

Bakhtiyar, Y., Arafat, M. Y., Andrabi, S., & Tak, H. I. (2020). Zooplankton: The Significant Ecosystem Service Provider in Aquatic Environment. In Bioremediation and Biotechnology, Vol 3: Persistent and Recalcitrant Toxic Substances (Vol. 3, pp. 227–244). Springer International Publishing. 10.1007/978-3-030-46075-4_10

Berga, M., Östman, Ö., Lindström, E. S., & Langenheder, S. (2015). Combined effects of zooplankton grazing and dispersal on the diversity and assembly mechanisms of bacterial metacommunities. Environmental Microbiology, 17(7), 2275–2287. 10.1111/1462-2920.12688

Bernardo-Cravo, A. P., Schmeller, D. S., Chatzinotas, A., Vredenburg, V. T., & Loyau, A. (2020). Environmental Factors and Host Microbiomes Shape Host–Pathogen Dynamics. In Trends in Parasitology (Vol. 36, Issue 7, pp. 616–633). Elsevier Ltd. 10.1016/j.pt.2020.04.010

Boscaro, V., Holt, C. C., Van Steenkiste, N. W. L., Herranz, M., Irwin, N. A. T., Àlvarez-Campos, P., Grzelak, K., Holovachov, O., Kerbl, A., Mathur, V., Okamoto, N., Piercey, R. S., Worsaae, K., Leander, B. S., & Keeling, P. J. (2022). Microbiomes of microscopic marine invertebrates do not reveal signatures of phylosymbiosis. Nature Microbiology, 7(6), 810–819. 10.1038/s41564-022-01125-9

Cáceres, C. E. (1998). Seasonal dynamics and interspecific competition in Oneida Lake *Daphnia*. Oecologia, 115, 233–244.

Cáceres, C. E., Tessier, A. J., Duffy, M. A., & Hall, S. R. (2014). Disease in freshwater zooplankton: What have we learned and where are we going. Journal of Plankton Research, 36(2), 326–333. 10.1093/plankt/fbt136

Callens, M., Watanabe, H., Kato, Y., Miura, J., & Decaestecker, E. (2018). Microbiota inoculum composition affects holobiont assembly and host growth in *Daphnia*. Microbiome, 6(1), 56. 10.1186/s40168-018-0444-1

Caporaso, J. G., Lauber, C. L., Walters, W. A., Berg-Lyons, D., Lozupone, C. A., Turnbaugh, P. J., Fierer, N., & Knight, R. (2011). Global patterns of 16S rRNA diversity at a depth of millions of sequences per sample. Proceedings of the National Academy of Sciences of the United States of America, 108(SUPPL. 1), 4516–4522. 10.1073/pnas.1000080107

Caughman, A. M., Pratte, Z. A., Patin, N. V, Frank, •, & Stewart, J. (2021). Coral microbiome changes over the day-night cycle. Coral Reefs, 40, 921–935. 10.1007/s00338

Cooper, R. O., Tjards, S., Rischling, J., Nguyen, D. T., & Cressler, C. E. (2022). Multiple generations of antibiotic exposure and isolation influence host fitness and the microbiome in a model zooplankton species. FEMS Microbiology Ecology, 98(10). 10.1093/femsec/fiac082

Decaestecker, E., Van de Moortel, B., Mukherjee, S., Gurung, A., Stoks, R., & De Meester, L. (2024). Hierarchical eco-evo dynamics mediated by the gut microbiome. In Trends in Ecology and Evolution (Vol. 39, Issue 2, pp. 165–174). Elsevier Ltd. 10.1016/j.tree.2023.09.013

De Corte, D., Varela, M. M., Louro, A. M., Bercovici, S. K., Valencia-Vila, J., Sintes, E., Baltar, F., Rodríguez-Ramos, T., Simon, M., Bode, A., Dittmar, T., & Niggemann, J. (2023). Zooplankton-derived dissolved organic matter composition and its bioavailability to natural prokaryotic communities. Limnology and Oceanography, 68(2), 336–347. 10.1002/lno.12272

Degans, H., & De Meester, L. (2002). Top-down control of natural phyto-and bacterioplankton prey communities by *Daphnia magna* and by the natural zooplankton community of the hypertrophic Lake Blankaart. In Hydrobiologia (Vol. 479).

Dixon, P. (2003). VEGAN, a package of R functions for community ecol. Journal of Vegetation Science, 14, 927–930.

Ebert, D. (2005). Ecology, Epidemiology and Evolution of Parasitism in Daphnia. Bethesda (MD) National Center for Biotechnology Information.

Eckert, E. M., Anicic, N., & Fontaneto, D. (2021). Freshwater zooplankton microbiome composition is highly flexible and strongly influenced by the environment. Molecular Ecology, 30(6), 1545–1558. 10.1111/mec.15815

Eckert, E. M., & Pernthaler, J. (2014). Bacterial epibionts of *Daphnia*: A potential route for the transfer of dissolved organic carbon in freshwater food webs. ISME Journal, 8(9), 1808–1819. 10.1038/ismej.2014.39

Edgar, R. C. (2016). SINTAX: a simple non-Bayesian taxonomy classifier for 16S and ITS sequences. BioRxiv. 10.1101/074161

Edgar, R. C., & Flyvbjerg, H. (2015). Error filtering, pair assembly and error correction for next-generation sequencing reads. Bioinformatics, 31(21), 3476–3482. 10.1093/bioinformatics/btv401

Epstein, H. E., Smith, H. A., Cantin, N. E., Mocellin, V. J. L., Torda, G., & van Oppen, M. J. H. (2019). Temporal variation in the microbiome of acropora coral species does not reflect seasonality. Frontiers in Microbiology, 10. 10.3389/fmicb.2019.01775

Escalas, A., Auguet, J. C., Avouac, A., Seguin, R., Gradel, A., Borrossi, L., & Villéger, S. (2021). Ecological Specialization Within a Carnivorous Fish Family Is Supported by a Herbivorous Microbiome Shaped by a Combination of Gut Traits and Specific Diet. Frontiers in Marine Science, 8. 10.3389/fmars.2021.622883

Frankel-Bricker, J., Song, M. J., Benner, M. J., & Schaack, S. (2020). Variation in the Microbiota Associated with *Daphnia magna* Across Genotypes, Populations, and Temperature. Microbial Ecology, 79(3), 731–742. 10.1007/s00248-019-01412-9

Freese, H. M., & Schink, B. (2011). Composition and Stability of the Microbial Community inside the Digestive Tract of the Aquatic Crustacean *Daphnia magna*. Microbial Ecology, 62(4), 882–894. 10.1007/s00248-011-9886-8

Frost, P. C., Ebert, D., & Smith, V. H. (2008). Bacterial infection changes the elemental composition of *Daphnia magna*. Journal of Animal Ecology, 77(6), 1265–1272. 10.1111/j.1365-2656.2008.01438.x

González-Tortuero, E., Rusek, J., Turko, P., Petrusek, A., Maayan, I., Piálek, L., Tellenbach, C., Gießler, S., Spaak, P., & Wolinska, J. (2016). *Daphnia* parasite dynamics across multiple *Caullerya* epidemics indicate selection against common parasite genotypes. Zoology, 119(4), 314–321. 10.1016/j.zool.2016.04.003

Gorokhova, E., El-Shehawy, R., Lehtiniemi, M., & Garbaras, A. (2021). How Copepods Can Eat Toxins Without Getting Sick: Gut Bacteria Help Zooplankton to Feed in Cyanobacteria Blooms. Frontiers in Microbiology, 11. 10.3389/fmicb.2020.589816

Hébert, M. P., Beisner, B. E., & Maranger, R. (2016). A meta-analysis of zooplankton functional traits influencing ecosystem function. Ecology, 97(4), 1069–1080. 10.1890/15-1084.1

Hegg, A., Radersma, R., & Uller, T. (2021). A field experiment reveals seasonal variation in the *Daphnia* gut microbiome. Oikos, 130(12), 2191–2201. 10.1111/oik.08530

Ichige, R., & Urabe, J. (2023). Divergence of the Host-Associated Microbiota with the Genetic Distance of Host Individuals Within a Parthenogenetic *Daphnia* Species. Microbial Ecology, 86(3), 2097–2108. 10.1007/s00248-023-02219-5

Infante-Villamil, S., Huerlimann, R., & Jerry, D. R. (2021). Microbiome diversity and dysbiosis in aquaculture. In Reviews in Aquaculture (Vol. 13, Issue 2, pp. 1077–1096). John Wiley and Sons Inc. 10.1111/raq.12513

Kimes, N. E., Johnson, W. R., Torralba, M., Nelson, K. E., Weil, E., & Morris, P. J. (2013). The *Montastraea faveolata* microbiome: Ecological and temporal influences on a Caribbean reef-building coral in decline. Environmental Microbiology, 15(7), 2082–2094. 10.1111/1462-2920.12130

Krajacich, B. J., Huestis, D. L., Dao, A., Yaro, A. S., Diallo, M., Krishna, A., Xu, J., & Lehmann, T. (2018). Investigation of the seasonal microbiome of *Anopheles coluzzii* mosquitoes in Mali. PLoS ONE, 13(3). 10.1371/journal.pone.0194899

Li, Y., Xu, Z., & Liu, H. (2021). Nutrient-imbalanced conditions shift the interplay between zooplankton and gut microbiota. BMC Genomics, 22(1). 10.1186/s12864-020-07333-z

Lohr, J. N., Laforsch, C., Koerner, H., & Wolinska, J. (2010). A *Daphnia* parasite (*Caullerya mesnili*) constitutes a new member of the ichthyosporea, a group of protists near the animal-fungi divergence. Journal of Eukaryotic Microbiology, 57(4), 328–336. 10.1111/j.1550-7408.2010.00479.x

López, M. F., Rebollar, E. A., Harris, R. N., Vredenburg, V. T., & Hero, J. M. (2017). Temporal variation of the skin bacterial community and *Batrachochytrium dendrobatidis* infection in the terrestrial cryptic frog *Philoria loveridgei*. Frontiers in Microbiology, 8(DEC). 10.3389/fmicb.2017.02535

Love, M. I., Huber, W., & Anders, S. (2014). Moderated estimation of fold change and dispersion for RNA-seq data with DESeq2. Genome Biology, 15(12). 10.1186/s13059-014-0550-8

Macke, E., Callens, M., De Meester, L., & Decaestecker, E. (2017). Host-genotype dependent gut microbiota drives zooplankton tolerance to toxic cyanobacteria. Nature Communications, 8(1). 10.1038/s41467-017-01714-x

Macke, E., Callens, M., Massol, F., Vanoverberghe, I., De Meester, L., & Decaestecker, E. (2020). Diet and Genotype of an Aquatic Invertebrate Affect the Composition of Free-Living Microbial Communities. Frontiers in Microbiology, 11. 10.3389/fmicb.2020.00380

McMurdie, P. J., & Holmes, S. (2013). Phyloseq: An R Package for Reproducible Interactive Analysis and Graphics of Microbiome Census Data. PLoS ONE, 8(4). 10.1371/journal.pone.0061217

Meyer, J. L., Paul, V. J., & Teplitski, M. (2014). Community shifts in the surface microbiomes of the coral *Porites astreoides* with unusual lesions. PLoS ONE, 9(6). 10.1371/journal.pone.0100316

Miller, E. T., Svanbäck, R., & Bohannan, B. J. M. (2018). Microbiomes as Metacommunities: Understanding Host-Associated Microbes through Metacommunity Ecology. In Trends in Ecology and Evolution (Vol. 33, Issue 12, pp. 926–935). Elsevier Ltd. 10.1016/j.tree.2018.09.002

Minich, J. J., Petrus, S., Michael, J. D., Michael, T. P., Knight, R., & Allen, E. E. (2020). Temporal, Environmental, and Biological Drivers of the Mucosal Microbiome in a Wild Marine Fish, Scomber japonicus. MSphere, 5(3). 10.1128/msphere.00401-20

Montero-Pau, J., Gómez, A., & Muñoz, J. (2008). Application of an inexpensive and high-throughput genomic DNA extraction method for the molecular ecology of zooplanktonic diapausing eggs. Limnology and Oceanography: Methods, 6(6), 218–222. 10.4319/lom.2008.6.218

Paix, B., van der Valk, E., & de Voogd, N. J. (2024). Dynamics, diversity, and roles of bacterial transmission modes during the first asexual life stages of the freshwater sponge *Spongilla lacustris*. Environmental Microbiome, 19(1). 10.1186/s40793-024-00580-7

Peerakietkhajorn, S., Tsukada, K., Kato, Y., Matsuura, T., & Watanabe, H. (2015). Symbiotic bacteria contribute to increasing the population size of a freshwater crustacean, *Daphnia magna*. Environmental Microbiology Reports, 7(2), 364–372. 10.1111/1758-2229.12260

Pérez-Pascual, D., Pérez-Cobas, A. E., Rigaudeau, D., Rochat, T., Bernardet, J. F., Skiba-Cassy, S., Marchand, Y., Duchaud, E., & Ghigo, J. M. (2021). Sustainable plant-based diets promote rainbow trout gut microbiota richness and do not alter resistance to bacterial infection. Animal Microbiome, 3(1). 10.1186/s42523-021-00107-2

Pfenning-Butterworth, A., Cooper, R. O., & Cressler, C. E. (2022). Daily feeding rhythm linked to microbiome composition in two zooplankton species. PLoS ONE, 17(2 February). 10.1371/journal.pone.0263538

Pierce, M. L., & Ward, J. E. (2019). Gut Microbiomes of the Eastern Oyster (*Crassostrea virginica*) and the Blue Mussel (*Mytilus edulis*): Temporal Variation and the Influence of Marine Aggregate-Associated Microbial Communities. MSphere, 4(6). 10.1128/msphere.00730-19

Porter, K. G., Gerritsen, J., & Orcutt, J. D. (1982). The effect of food concentration on swimming patterns, feeding behavior, ingestion, assimilation, and respiration by Daphnia. Limnology and Oceanography, 27(5), 935–949. 10.4319/lo.1982.27.5.0935

Preston, D. L., Mischler, J. A., Townsend, A. R., & Johnson, P. T. J. (2016). Disease Ecology Meets Ecosystem Science. Ecosystems, 19(4), 737–748. 10.1007/s10021-016-9965-2

Quast, C., Pruesse, E., Yilmaz, P., Gerken, J., Schweer, T., Yarza, P., Peplies, J., & Glöckner, F. O. (2013). The SILVA ribosomal RNA gene database project: Improved data processing and web-based tools. Nucleic Acids Research, 41(D1). 10.1093/nar/gks1219

Rajarajan, A., Wolinska, J., Walser, J. C., Dennis, S. R., & Spaak, P. (2023). Host-Associated Bacterial Communities Vary Between *Daphnia galeata* Genotypes but Not by Host Genetic Distance. Microbial Ecology, 85(4), 1578–1589. 10.1007/s00248-022-02011-x

Rajarajan, A., Wolinska, J., Walser, J. C., Mäder, M., & Spaak, P. (2022). Infection by a eukaryotic gut parasite in wild *Daphnia sp.* associates with a distinct bacterial community. FEMS Microbiology Ecology, 98(10). 10.1093/femsec/fiac097

R Core Team. (2023). R: A Language and Environment for Statistical Computing. https://www.R-project.org/

Russell, S. L. (2019). Transmission mode is associated with environment type and taxa across bacteria-eukaryote symbioses: A systematic review and meta-analysis. In FEMS Microbiology Letters (Vol. 366, Issue 3). Oxford University Press. 10.1093/femsle/fnz013

Schmeller, D. S., Cheng, T., Shelton, J., Lin, C. F., Chan-Alvarado, A., Bernardo-Cravo, A., Zoccarato, L., Ding, T. S., Lin, Y. P., Swei, A., Fisher, M. C., Vredenburg, V. T., & Loyau, A. (2022). Environment is associated with chytrid infection and skin microbiome richness on an amphibian rich island (Taiwan). Scientific Reports, 12(1). 10.1038/s41598-022-20547-3

Schoebel, C. N., Tellenbach, C., Spaak, P., & Wolinska, J. (2011). Temperature effects on parasite prevalence in a natural hybrid complex. Biology Letters, 7(1), 108–111. 10.1098/rsbl.2010.0616

Sehnal, L., Brammer-Robbins, E., Wormington, A. M., Blaha, L., Bisesi, J., Larkin, I., Martyniuk, C. J., Simonin, M., & Adamovsky, O. (2021). Microbiome Composition and Function in Aquatic Vertebrates: Small Organisms Making Big Impacts on Aquatic Animal Health. In Frontiers in Microbiology (Vol. 12). Frontiers Media S.A. 10.3389/fmicb.2021.567408

Sharp, K. H., Pratte, Z. A., Kerwin, A. H., Rotjan, R. D., & Stewart, F. J. (2017). Season, but not symbiont state, drives microbiome structure in the temperate coral *Astrangia poculata*. Microbiome, 5(1), 120. 10.1186/s40168-017-0329-8

Shoemaker, K. M., Duhamel, S., & Moisander, P. H. (2019). Copepods promote bacterial community changes in surrounding seawater through farming and nutrient enrichment. Environmental Microbiology, 21(10), 3737–3750. 10.1111/1462-2920.14723

Sison-Mangus, M. P., Mushegian, A. A., & Ebert, D. (2015). Water fleas require microbiota for survival, growth and reproduction. ISME Journal, 9(1), 59–67. 10.1038/ismej.2014.116

Stencel, A. (2021). Do seasonal microbiome changes affect infection susceptibility, contributing to seasonal disease outbreaks? BioEssays, 43(1). 10.1002/bies.202000148

Stillson, P. T., Baltrus, D. A., & Ravenscraft, A. (2022). Prevalence of an Insect-Associated Genomic Region in Environmentally Acquired *Burkholderiaceae* Symbionts. Applied and Environmental Microbiology, 88(9). 10.1128/aem.02502-21

Sullam, K. E., Pichon, S., Schaer, T. M. M., & Ebert, D. (2018). The Combined Effect of Temperature and Host Clonal Line on the Microbiota of a Planktonic Crustacean. Microbial Ecology, 76(2), 506–517. 10.1007/s00248-017-1126-4

Sze, M., Doonan, J., McDonald, J., & Dewar, M. (2020). Factors that shape the host microbiome. In R. E. Antwis, X. A. Harrison, & M. J. Cox (Eds.), Microbiomes of Soils, Plants and Animals. Cambridge University Press. 10.1017/9781108654418

Talling, J. F. (2003). Phytoplankton-zooplankton seasonal timing and the “clear-water phase” in some English lakes. Freshwater Biology, 48(1), 39–52. 10.1046/j.1365-2427.2003.00968.x

Tang, K. W. (2005). Copepods as microbial hotspots in the ocean: effects of host feeding activities on attached bacteria. Aquatic Microbial Ecology, 38, 31–40.

Tellenbach, C., Tardent, N., Pomati, F., Keller, B., Hairston, N. G., Wolinska, J., & Spaak, P. (2016). Cyanobacteria facilitate parasite epidemics in *Daphnia*. Ecology, 97(12), 3422–3432. 10.1002/ecy.1576

Turko, P., Tellenbach, C., Keller, E., Tardent, N., Keller, B., Spaak, P., & Wolinska, J. (2018). Parasites driving host diversity: Incidence of disease correlated with *Daphnia* clonal turnover. Evolution, 72(3), 619–629. 10.1111/evo.13413

Utermöhl, H. (1958). Zur Vervollkommung der quantitativen phytoplankton-methodik.

Wagner, A. (2008). Light limitation increases the edibility of *Asterionella formosa* Hass. for *Daphnia* during periods of ice cover. Limnologica, 38(3–4), 286–301. 10.1016/j.limno.2008.06.004

Wang, Y., Smith, H. K., Goossens, E., Hertzog, L., Bletz, M. C., Bonte, D., Verheyen, K., Lens, L., Vences, M., Pasmans, F., & Martel, A. (2021). Diet diversity and environment determine the intestinal microbiome and bacterial pathogen load of fire salamanders. Scientific Reports, 11(1). 10.1038/s41598-021-98995-6

Wolinska, J., Keller, B., Manca, M., & Spaak, P. (2007). Parasite Survey of a *Daphnia* Hybrid Complex: Host-Specificity and Environment. Journal of Animal Ecology, 76(1), 191–200. https://www.jstor.org/stable/4125109

Xu, T., Novotny, A., Zamora-Terol, S., Hambäck, P. A., & Winder, M. (2024). Dynamics of Gut Bacteria Across Different Zooplankton Genera in the Baltic Sea. Microbial Ecology, 87(1). 10.1007/s00248-024-02362-7

Yoshida, T., Kagami, M., Gurung, T. B., & Urabe, J. (2001). Seasonal succession of zooplankton in the north basin of Lake Biwa. In Aquatic Ecology (Vol. 35).

Zhu, J., Li, H., Jing, Z. Z., Zheng, W., Luo, Y. R., Chen, S. X., & Guo, F. (2022). Robust host source tracking building on the divergent and non-stochastic assembly of gut microbiomes in wild and farmed large yellow croaker. Microbiome, 10(1). 10.1186/s40168-021-01214-7

Ziegler, M., Grupstra, C. G. B., Barreto, M. M., Eaton, M., BaOmar, J., Zubier, K., Al-Sofyani, A., Turki, A. J., Ormond, R., & Voolstra, C. R. (2019). Coral bacterial community structure responds to environmental change in a host-specific manner. Nature Communications, 10(1). 10.1038/s41467-019-10969-5

