## Supplementary material for "*Daphnia*-associated bacterial communities correlate with diet quantity, environmental conditions and epidemic size across natural outbreaks"

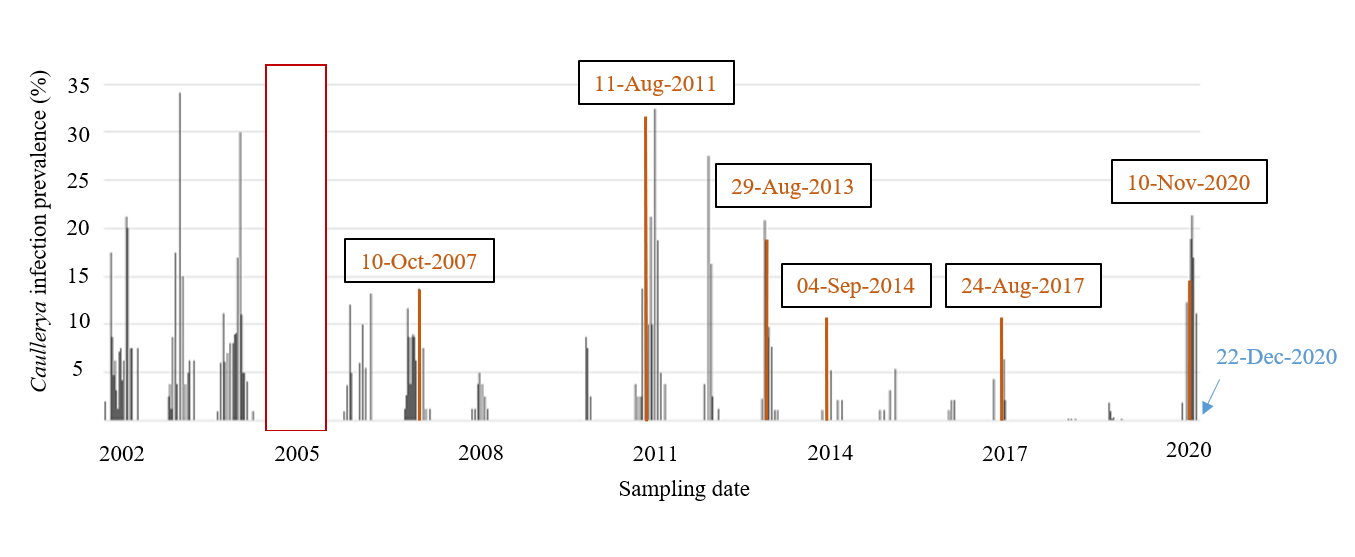
**Fig. S1** Prevalence of *Caullerya* infection (epidemic size) among *Daphnia* sp. in Lake Greifensee over time. Orange bars, marked with dates, indicate sampling events during which infected and uninfected *Daphnia* were collected for the present study. The blue arrow points to the sampling event from (Rajarajan, Wolinska et al. 2022). No data were collected during 2005 (red box).

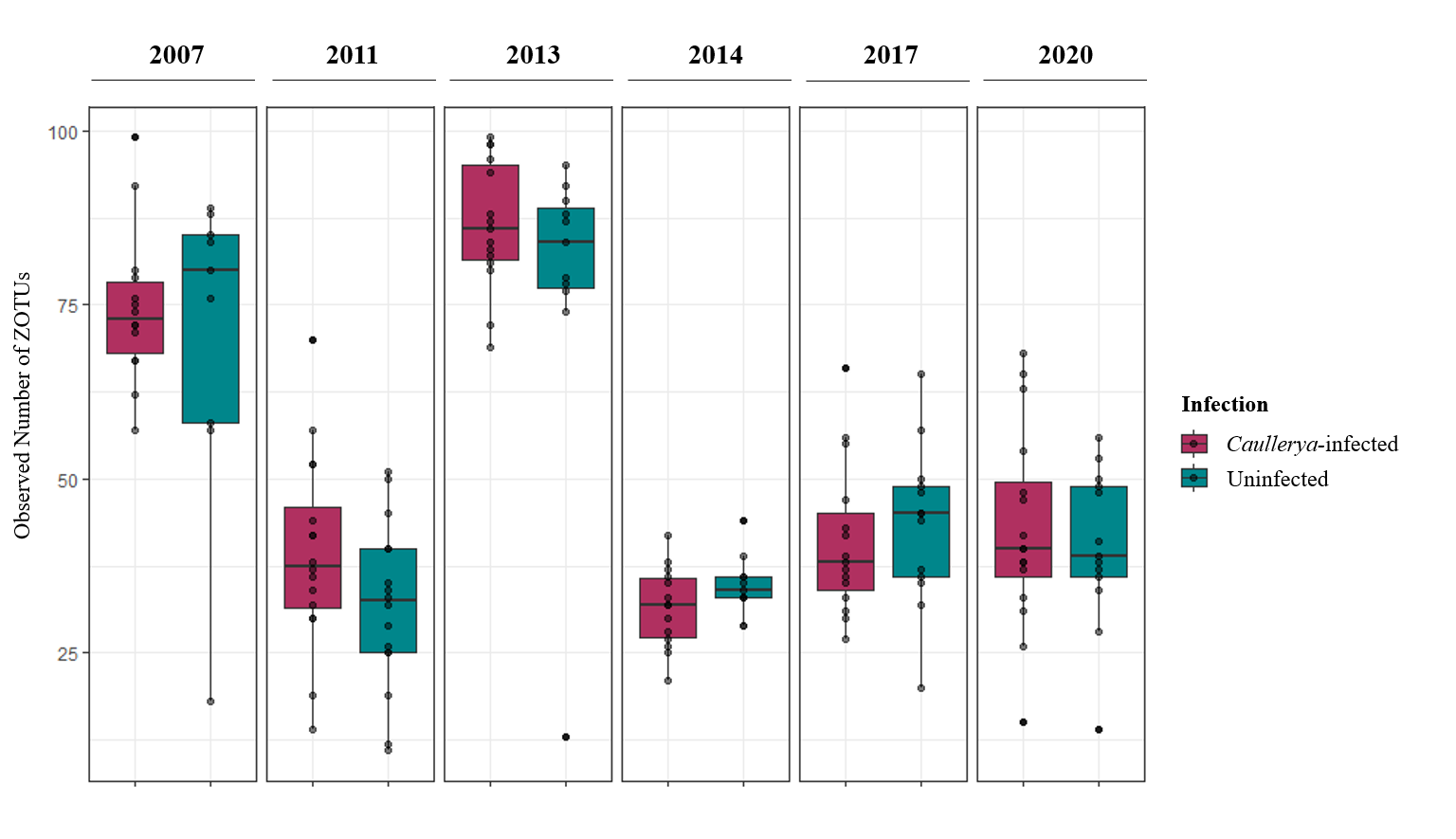

**Fig. S2** Alpha diversity (observed ZOTU richness) of samples over time and by infection status. Older samples were frozen for longer periods before sequencing, but the observed number of ZOTUs shows no clear trend of increase or decrease over time. In general, long-term freezing at -80°C is supposed to have a negligible effect on the sequenced bacterial communities (Tap, Cools-Portier et al. 2019).

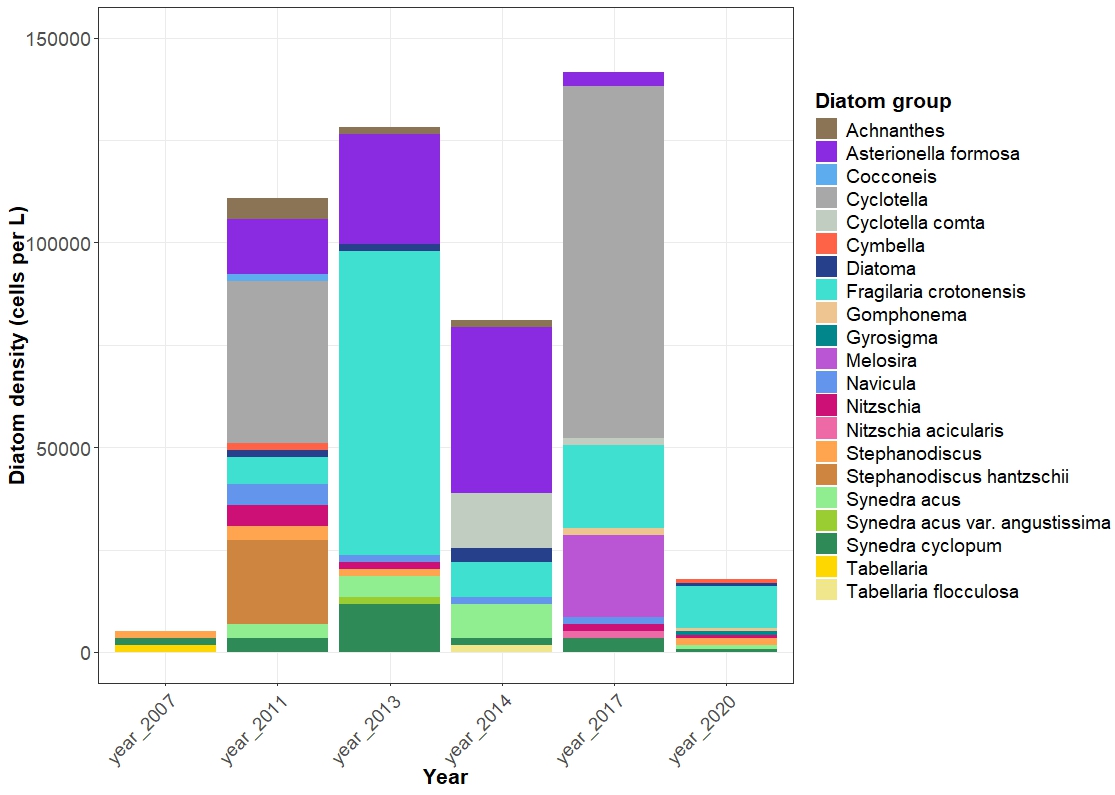

**Fig. S3** Density (cells L^-1^) of diatom groups in lake water during each year, displayed with their lowest identified taxonomic level.

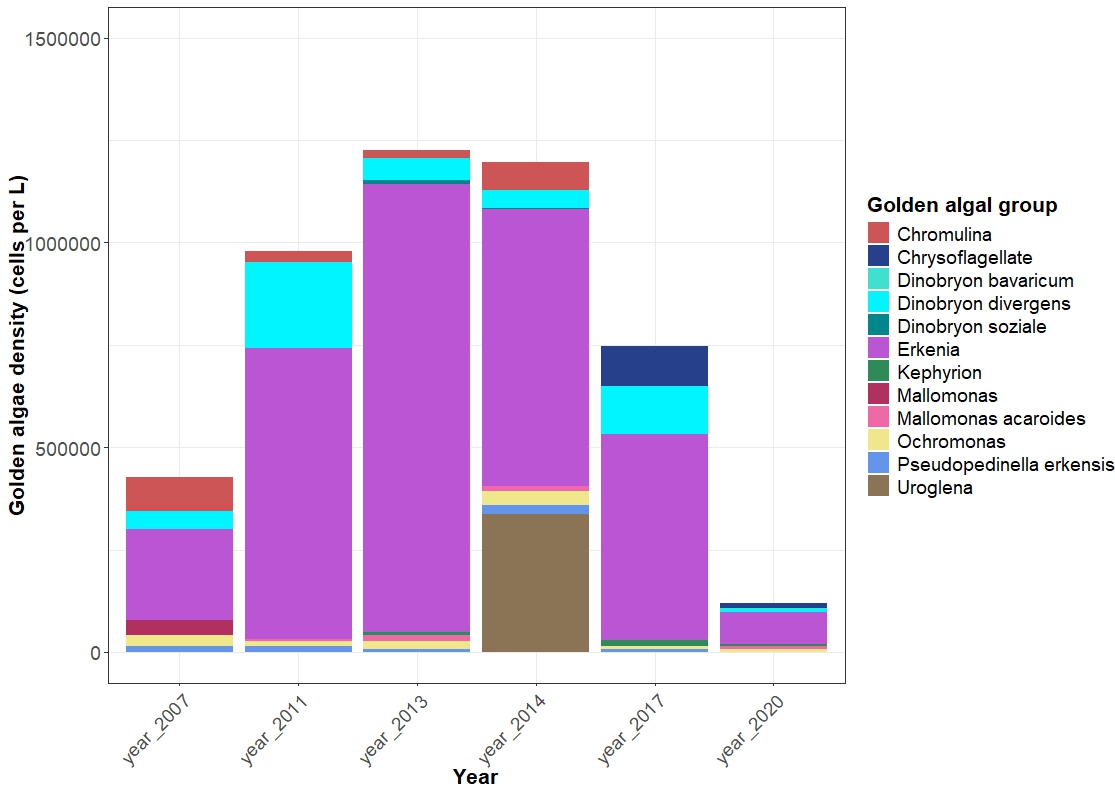

**Fig. S4** Density (cells L^-1^) of golden algal groups in lake water during each year, displayed with their lowest identified taxonomic level. *Erkenia* (=*Chrysochromulina* sp.) dominated all years.

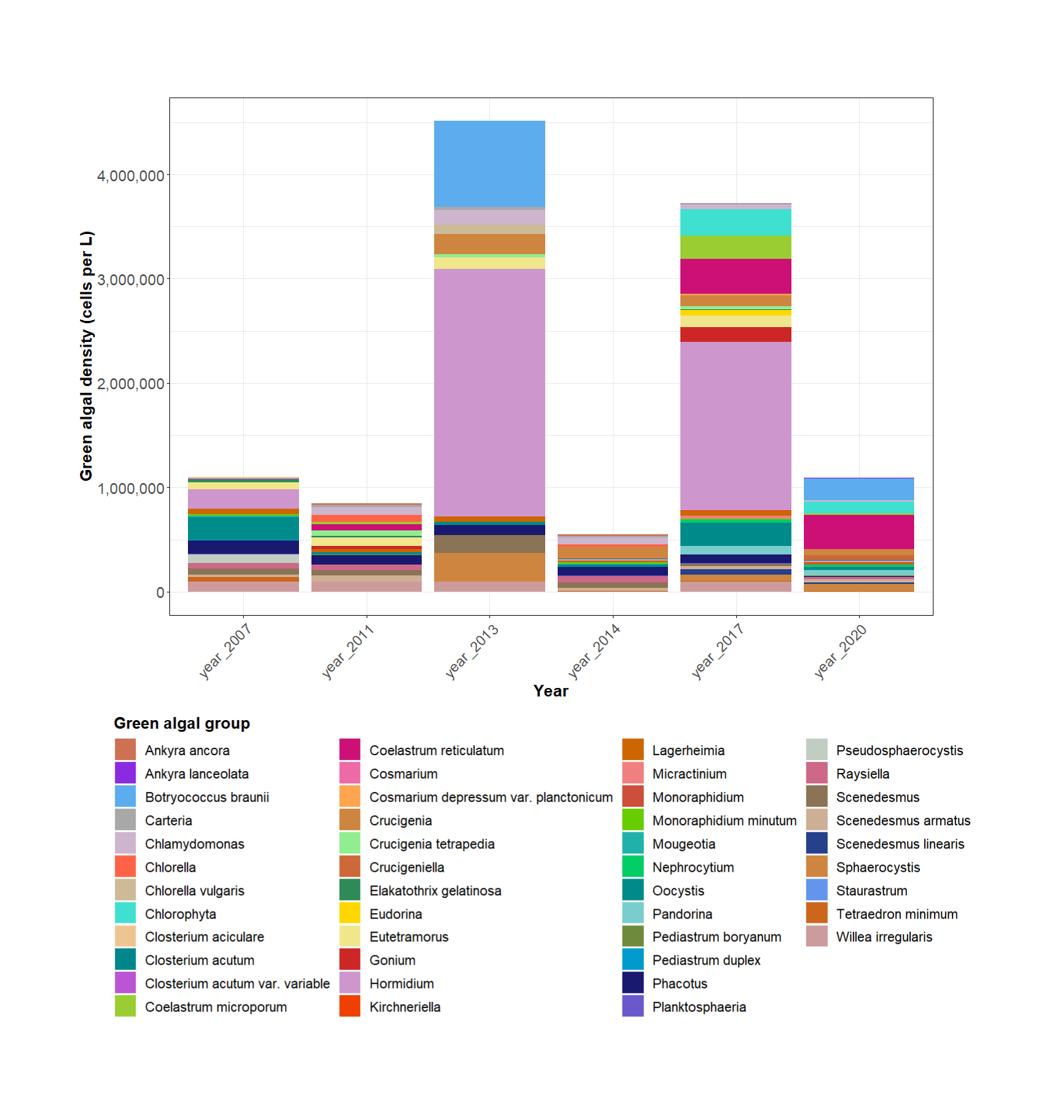

**Fig. S5** Density (cells L^-1^) of green algal groups in lake water during each year, displayed with their lowest identified taxonomic level.

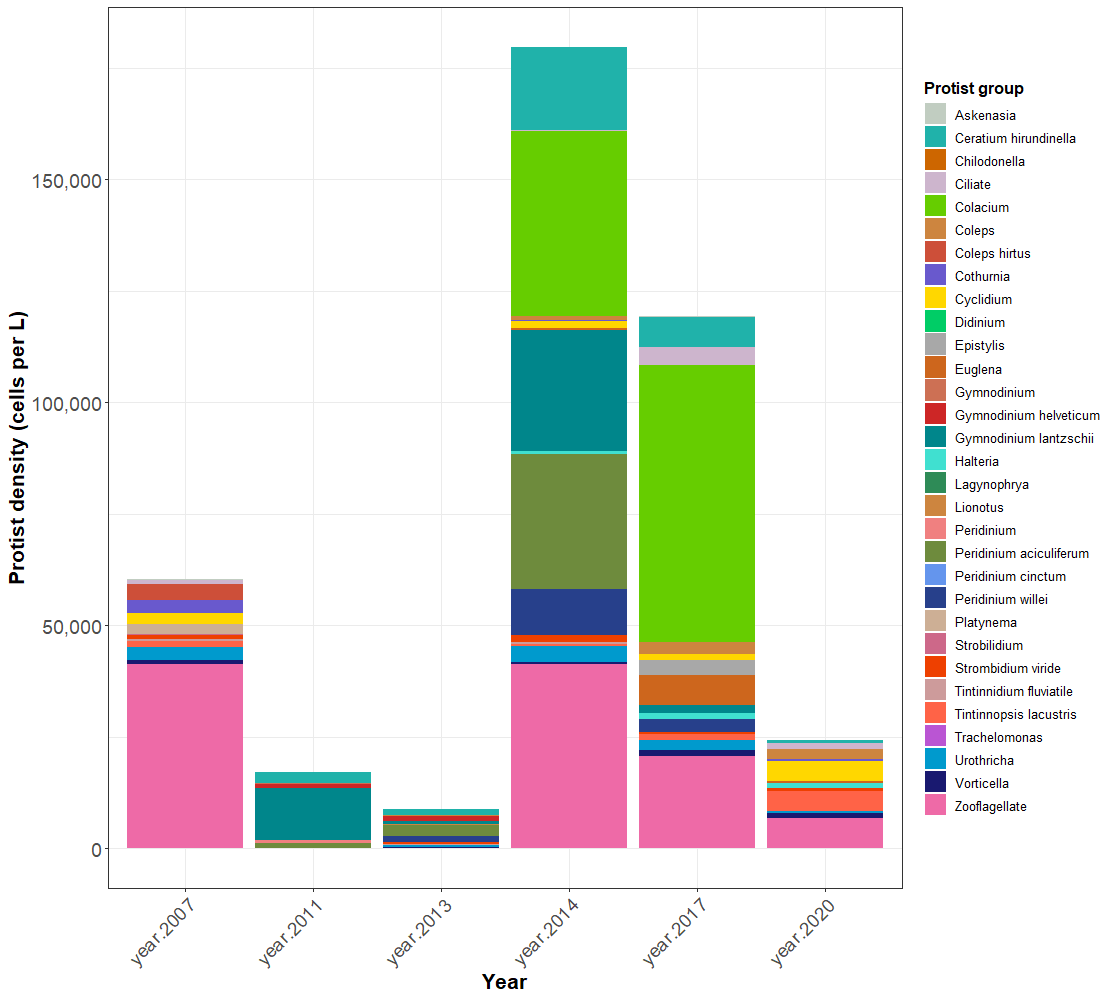

**Fig. S6** Density (cells L^-1^) of protist groups in lake water during each year, displayed with their lowest identified taxonomic level.

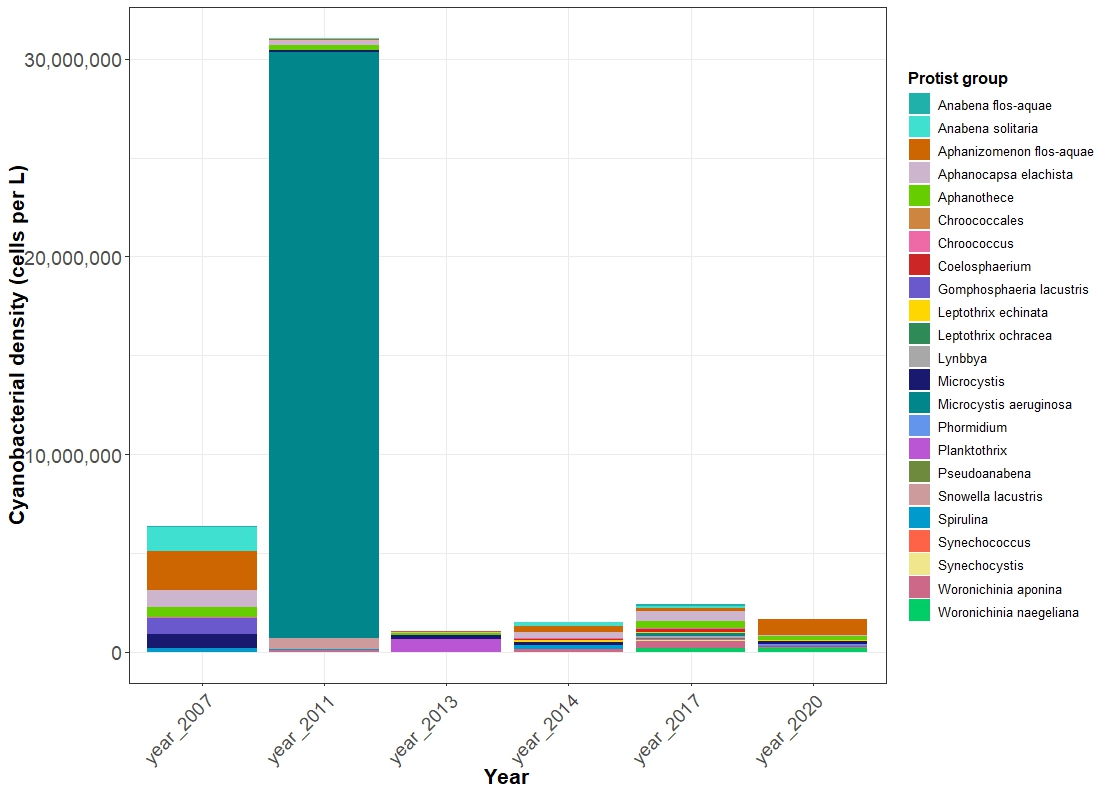

**Fig. S7** Density (cells L^-1^) of cyanobacterial groups in lake water during each year, displayed with their lowest identified taxonomic level.

| **Year** | ***Daphnia* sampling date** | **Date of physicochemical measurement** | **Date of phyto- and zooplankton data collection** | **# *Caullerya-*infected *Daphnia*** | **# Uninfected *Daphnia*** |
| --- | --- | --- | --- | --- | --- |
| 2007 | 10.10.2007 | 08.10.2007 | 08.10.2007 | 14 | 9 |
| 2011 | 11.08.2011 | 15.08.2011 | 16.08.2011 | 16 | 16 |
| 2013 | 29.08.2013 | 02.09.2013 | 03.09.2013 | 15 | 11 |
| 2014 | 04.09.2014 | 01.09.2014 | 04.09.2014 | 14 | 11 |
| 2017 | 24.08.2017 | 04.09.2017 | 05.09.2017 | 15 | 13 |
| 2020 | 10.11.2020 | 02.11.2020 | 04.11.2020 | 16 | 13 |

**Table S1** Dates for (a) sampling events of Daphnia, (b) abiotic data collection (temperature and dissolved oxygen), and (c) phytoplankton and zooplankton data (density and community composition). The “Caullerya-infected” and “Uninfected” columns state the final sample sizes of Daphnia analyzed in the present study.
